# Nanoparticle-mediated transformation expands horizon of organism engineering

**DOI:** 10.1101/559252

**Authors:** Katherine E. French

## Abstract

Our ability to engineer organisms will unlock new discoveries in how cellular processes work and enable new advances in biotechnology. However, many organisms are recalcitrant to traditional transformation methods. Here, we describe a new nanoparticle-based method of transformation which can be used to transform bacteria, plants, algae and diatoms.

---

Our ability to engineer organisms will unlock new discoveries in how cellular processes work and enable new advances in biotechnology (Khalil and Collins 2010). This process is known as ‘transformation’ and relies on electric currents, heat shock, or chemicals to create tiny pores in organism cell membranes that allow foreign genetic material to enter the host. However, many organisms are recalcitrant to these methods (Brenner et al. 2008). Recent advances in nanoparticle technology have illustrated that quantum dots can deliver siRNA (typically of 20-22 bp) and other molecules to human and mice cells, but these methods do not work on bacteria or organisms with cell walls (Kim et al. 2017; Pierrat et al. 2015). Here, we describe a new carbon-nanoparticle based transformation system. Nanoparticle composition, polymer coating, pH, and plasmid DNA (p-DNA) concentration were key factors in transformation success. Pilot experiments with eukaryotic, prokaryotic, and difficult-to-transform species suggest any organism that can take-up the nanoparticles we designed can be transformed using this method.

We prepared 15 different nanoparticles to assess the following properties (1) organism uptake; (2) cellular localization; (3) transformation ability; and (4) cellular toxicity. We replicated two nanoparticle types from previous studies; the rest of the nanoparticles are unique to this study. For nanoparticles with desired optical and functional properties, we experimented with different polymer coatings (bPEI and PEG), polymer concentrations (0.0125-0.05 g/mL), plasmid concentrations (1 ng-1 ug/ul), and nanoparticle-plasmid incubation time (10 min. to 24 h). The particles best suited for general (non-organelle) transformation were made of citric acid or glucose, passivated with low concentrations of bPEI (0.01-0.02 g/mL) and plasmid (1-4ng/ul), and with a short nanoparticle-plasmid conjugation time (15-20 min.). Particles made with threonine localized to chloroplasts and those made with arginine, lysine, valine and serine localized to mitochondria (**SI Table 1**). Nanoparticles made with specific amino acid combinations may mimic the role of terminal sequences in guiding proteins targeted to specific cellular compartments (Carrie et al. 2009; de Castro et al. 1996; Lodish et al. 2000).

Cellular uptake studies were carried out in the following organisms: plants/mosses (*Arabidopsis thaliana, Nicotiana benthamiana, Physcomitrella patens* and *Selaginella kraussiana*), protists (*Physarum polycephalum*), bacteria (*Pseudomonas flourescens*), cyanobacteria (*Synecoccochus*), diatoms (*Thalassiosira oceanica*), algae (*Penium margaritaceum, Spyrogira* sp.), fungi (*Trichoderma reesei*), and coccolithophores (*Pleurochrysis certerae, Gephyrocapsa oceanica*). This is the first study to show algae, diatoms and coccolithophores will take up nanoparticles. Nanoparticles entered cells in two ways: (1) through membrane pores or (2) invagination of cell membranes. For example, in diatoms nanoparticles appear to enter the cell through external pores in the frustule; in algae such as *Penium*, in-folding of the cell membrane around nanoparticle aggregates was observed and these vesicles seemed to transport nanoparticles in and out of the cell (**Fig. 1 A and B**). The latter correlates with Giraldo et al. 2014, who proposed that single-walled carbon nanotubes may enter the double-membranes of chloroplasts via diffusion, becoming coated with glycerolipids in the process. Experiments with vascular plants, filamentous fungi, and *P. polycephalum* indicate nanoparticles can be transported long distances (either within the vascular system or within vesicles) (**SI Fig. 1**). Based on material composition, nanoparticles localized to six places: the cell membrane, cytoplasm, vacuole, nucleus, mitochondria, and chloroplasts.

**Fig. 1.**
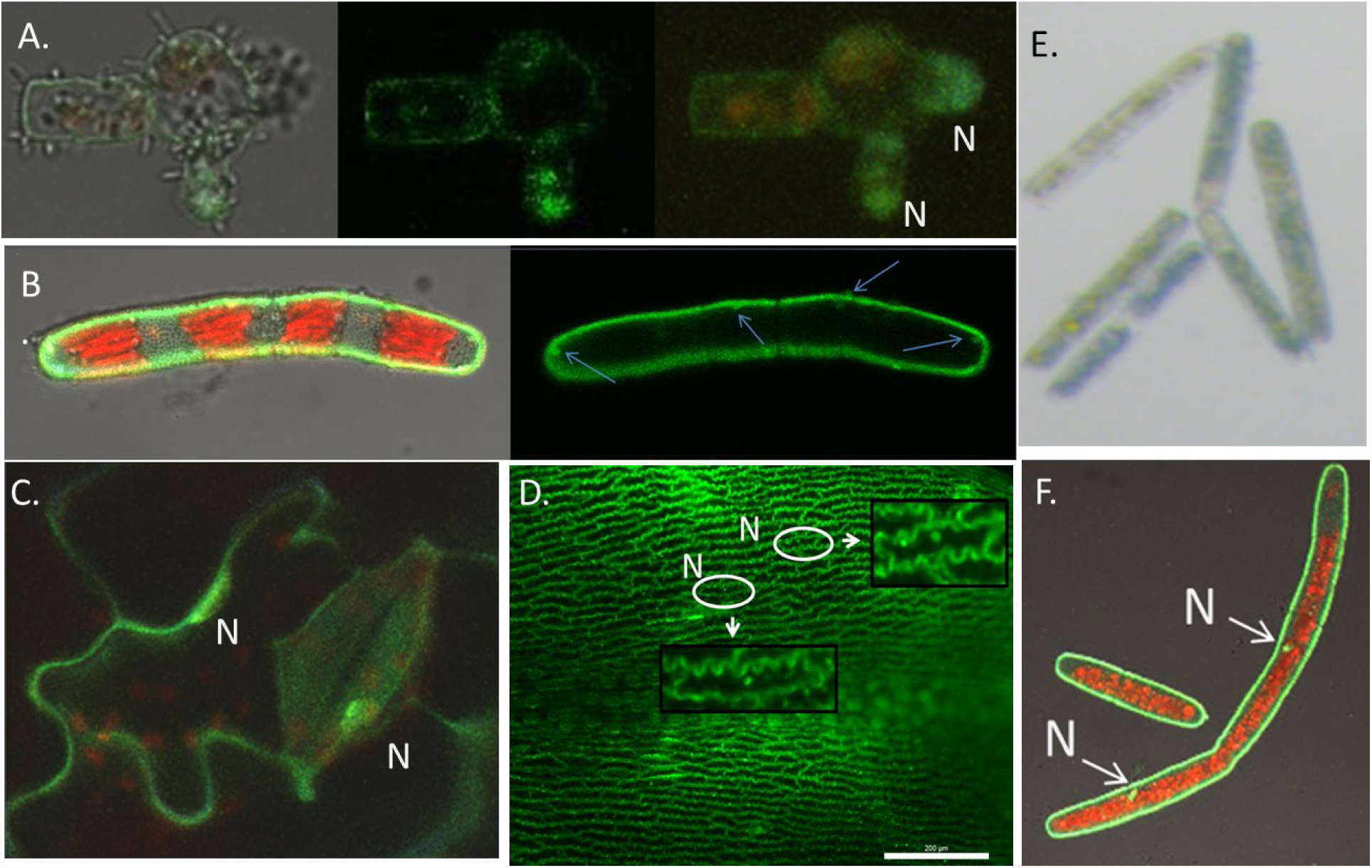
Organism uptake of carbon nanoparticles and transformation. A. Expression on VNLS in *T. oceanica*. The image shows dorsal and lateral views of several diatoms in different stages of life. B. Uptake of CNDs in *P. margaritaceum*. Arrows point to vesicular structures filled with nanoparticles. C. Transformation of *N. benthamiana* with VNLS. D. Expression of VNLS in *S. kraussiana* 16 hours after exposure to CND-pDNA complexes; boxes show magnified cells with fluorescent nuclei. E. GUS-staining of *P. margaritaceum*. F. Dividing *P. margaritaceum* eight days after exposure to CND-pDNA complexes. N= nucleus. Confocal images (A, B, C, F) taken using excitation 405/488/543nm. Images are composed of superimposed z-sections. Images D and E were taken with a Leica light microscope using a YFP filter and an LED light respectively.

In general, carbon-based nanoparticles are non-toxic to cells. This is in contrast to nanoparticles made out of heavy metals (e.g. ‘quantum dots’) which can kill cells and may cause environmental harm (Hardman 2006; Valizadeh et al. 2012). Study organisms did show signs of stress when exposed to nanoparticles with high bPEI (0.05-0.1 g/mL) and plasmid concentrations (300 ng-1 ug/ul) or with low pH (< pH 4). Signs of stress included retraction of the cytoplasm from the cell wall in vascular plants; cell rupture; and loss of innate chloroplast structure (specifically in algae). Once nanoparticles were optimized to decrease cell toxicity, organisms could survive on media (agar gel plates or liquid growth media) containing nanoparticles for over 30 days.

The following organisms were successfully transformed using glucose and citric acid based nanoparticles coated with bPEI and conjugated with a Venus NLS plasmid, a MCherry NLS plasmid, or a GFP HDEL plasmid: *A. thaliana, N. benthamiana, S. kraussiana, P. margaritaceum* and *T. oceanica* (**Fig. 1 A, C-F**; **SI Fig. 2**). Organisms were exposed to the following treatments: control (no treatment), bPEI, bPEI + pDNA, CND, CND + pDNA. No transformation was seen using plasmids bound to bPEI, a transfection method commonly used in mammalian cells (Kim et al. 2017; Pierrat et al. 2015), and exposure to bPEI alone resulted in cell death for plants and algae. Successful transformation using plasmids bound to CNDs was indicated by fluorescence of the nuclear envelope and the endoplasmic reticulum. To further confirm transformation, we transformed *P. margaritaceum* with a 35S ubi-GUS plasmid. Non-transformed cells appear clear or yellowish; transformed cells appear blue (**Fig. 1 E**). This system can be used for transient and stable expression of foreign genes in both chassis organisms and difficult to transform species. For example, the alga *P. margaritaceum* showed nuclear YFP and MCherry labelling eight days after transfection, suggesting the plasmid DNA was inserted into the hosts’ genomes. Other images show cells dividing, with both cells exhibiting nuclear fluorescence, suggesting the DNA is inherited (**Fig. 1 F**).

Carbon nanoparticle-based transformation presents a promising new method of altering the genetic make-up of cells. This is the first study to show the transport and expression of large plasmids (>10 kbp) into recalcitrant organisms using carbon nanoparticles. Future studies should look at the metabolic impact of CND exposure, the degradation of CNDs in cells and the environment, plasmid size limitations which may affect transformation, and whether organelle-targeting nanoparticles can be used to genetically modify chloroplasts and mitochondria. Further development of this technology will lead to new discoveries across the life sciences and commercial gains in the growing biotech sector.

## Author contributions

KEF designed the nanoparticles, performed the nanoparticle-uptake and transformation experiments, imaged the experiments, and wrote the manuscript.

