## Supplementary material for "Nanoparticle-mediated transformation expands horizon of organism engineering": SI

Katherine E. French*[1]

[1] Department of Plant and Microbiology, University of California Berkeley, 111 Koshland Hall, Berkeley, CA

**SI Methods**

1. Nanoparticle creation

We modified the protocols for citric acid and urea and citric acid and di-ammonium hydrogen phosphate from previous studies and created de novo protocols for the production of nanoparticles for transformation and organelle targeting. All reagents used were from Sigma. The protocols for each are described below. All nanoparticle solutions were stored at room temperature.

1.1. **Citric acid and urea**: We modified the protocol from Qu et al. (2012) to produce nanoparticles with green, turquoise, yellow and green emission spectra. We added 3g citric acid and 3g urea to 10 mL of dH20 in a glass beaker. After mixing the contents well, we microwaved the solution in a 750 watt oven for 4-5 minutes in a 50mL glass beaker. To break down the resulting carbon mass, we added 20 mL of dH20 and pulverized the carbonized material with a metal spatula. The mixture was poured into a 50 mL falcon tube and centrifuged at 15000 rpm for 20 min. The supernatant was collected with a 10 mL syringe and filtered with a 0.45 µm Millipore filter. The solution was purified using a Sephadex column (G100) connected to an automatic fraction collector. 500 ul fractions were collected and grouped by emission spectra (e.g. blue, turquoise, green, yellow).

1.2. **Citric acid and di-ammonium hydrogen phosphate**: We modified the protocol from Chandra et al. (2016) to produce nanoparticles doped with phosphorus. We mixed 0.3 g citric acid and 0.754g di-ammonium hydrogen phosphate with 20 mL dH20 in a beaker. We transferred the solution to a glass tube and covered with a heat-resistant rubber cover. The glass tube was place in a rack in an oven set at 180 C for 4 h. After cooling, we broke down the carbonized material with a metal spatula and added 20mL dH20 to re-suspend the carbonized material. We poured the liquid into a 50 mL falcon tube and centrifuged it at 15000 rpm for 20 min. The supernatant was collected with a 10 mL syringe and filtered with a 0.45 µm Millipore filter.

**General transformation**

**1.2 Citric acid and bpei**

We added 1 or 0.5g citric acid and 0.2, 0.1 or 0.5g bPEI 25k (made from the 100mg stock solution of bPEI listed in the notes) to 5 mL of 0.1 M HCL in a beaker and mixed the contents well, adding dH20 to bring the total volume of the reaction to 10mL. We then microwaved the solution in a 750 watt oven for 120s. After removing the beaker from the microwave, we broke down the carbonized mass with a spatula. We added 15 mL of dH20 and continued to break down material in beaker. We then poured the liquid into a 50 mL falcon tube. To remove large particles, we centrifuged the solution at 15000 rpm for 20 min. We then removed the liquid supernatant with a 20 mL syringe and filtered with a 0.22 µm Millipore filter. To decrease the acidity of the nanoparticle solution, we changed the pH of the solution to 6 with KOH. To remove any large particles created by the pH change, we centrifuged the solution at 15000 rpm for 20 minutes, removed the supernatant with a 20 mL syringe, and filtered the solution with a 0.22 µm Millipore filter. We stored the particles at room temperature.

**1.3 Glucose and bpei**

We added 1 or 0.5g glucose and 0.2, 0.1 or 0.5g bPEI 25k (made from the 100mg stock solution of bPEI listed in the notes) to 5 mL of 0.1 M HCL in a beaker and mixed the contents well, adding dH20 to bring the total volume of the reaction to 10mL. We then microwaved the solution in a 750 watt oven for 120s. After removing the beaker from the microwave, we broke down the carbonized mass with a spatula. We added 15 mL of dH20 and continued to break down material in beaker. We then poured the liquid into a 50 mL falcon tube. To remove large particles, we centrifuged the solution at 15000 rpm for 20 min. We then removed the liquid supernatant with a 20 mL syringe and filtered with a 0.22 µm Millipore filter. To decrease the acidity of the nanoparticle solution, we changed the pH of the solution to 6 with KOH. To remove any large particles created by the pH change, we centrifuged the solution at 15000 rpm for 20 minutes, removed the supernatant with a 20 mL syringe, and filtered the solution with a 0.22 µm Millipore filter. We stored the particles at room temperature.

**1.4 Dextrin and bpei**

We added 1 or 0.5g dextrin and 0.2, 0.1 or 0.5g bPEI 25k (made from the 100mg stock solution of bPEI listed in the notes) to 5 mL of 0.1 M HCL in a beaker and mixed the contents well, adding dH20 to bring the total volume of the reaction to 10mL. We then microwaved the solution in a 750 watt oven for 120s. After removing the beaker from the microwave, we broke down the carbonized mass with a spatula. We added 15 mL of dH20 and continued to break down material in beaker. We then poured the liquid into a 50 mL falcon tube. To remove large particles, we centrifuged the solution at 15000 rpm for 20 min. We then removed the liquid supernatant with a 20 mL syringe and filtered with a 0.22 µm Millipore filter. To decrease the acidity of the nanoparticle solution, we changed the pH of the solution to 6 with KOH. To remove any large particles created by the pH change, we centrifuged the solution at 15000 rpm for 20 minutes, removed the supernatant with a 20 mL syringe, and filtered the solution with a 0.22 µm Millipore filter. We stored the particles at room temperature.

**Organelle targeting**

**Nucleus-targeting**

**1.5 Citric acid and Lysine**

We added 3g citric acid, 3g lysine and 0.1 or 0.05g bPEI 25k (made from the 100mg stock solution of bPEI listed in the notes) to 5 mL of 0.1 M HCL in a beaker and mixed the contents well, adding dH20 to bring the total volume of the reaction to 10mL. We then microwaved the solution in a 750 watt oven for 120s. A yellow liquid will form. After removing the beaker from the microwave, we broke down the carbonized mass with a spatula. We added 15 mL of dH20 and continued to break down material in beaker. We then poured the liquid into a 50 mL falcon tube. To remove large particles, we centrifuged the solution at 15000 rpm for 20 min. We then removed the liquid supernatant with a 20 mL syringe and filtered with a 0.22 µm Millipore filter. To decrease the acidity of the nanoparticle solution, we changed the pH of the solution to 6 with KOH. To remove any large particles created by the pH change, we centrifuged the solution at 15000 rpm for 20 minutes, removed the supernatant with a 20 mL syringe, and filtered the solution with a 0.22 µm Millipore filter. We stored the particles at room temperature.

**1.6 Citric acid and Arginine**

We added 3g citric acid, 3g Arginine and 0.1 or 0.05g bPEI 25k (made from the 100mg stock solution of bPEI listed in the notes) to 5 mL of 0.1 M HCL in a beaker and mixed the contents well, adding dH20 to bring the total volume of the reaction to 10mL. We then microwaved the solution in a 750 watt oven for 120s. A yellow, sticky mass will form. After removing the beaker from the microwave, we broak down the carbonized mass with a spatula. We added 20 mL of dH20 and continued to break down material in beaker. We then poured the liquid into a 50 mL falcon tube. To remove large particles, we centrifuged the solution at 15000 rpm for 20 min. We then removed the liquid supernatant with a 20 mL syringe and filtered with a 0.22 µm Millipore filter. To decrease the acidity of the nanoparticle solution, we changed the pH of the solution to 6 with KOH. To remove any large particles created by the pH change, we centrifuged the solution at 15000 rpm for 20 minutes, removed the supernatant with a 20 mL syringe, and filtered the solution with a 0.22 µm Millipore filter. We stored the particles at room temperature.

**1.7 Glucose and Arginine**

We added 3g glucose, 3g arginine and 0.1 or 0.05g bPEI 25k (made from the 100mg stock solution of bPEI listed in the notes) to 5 mL of 0.1 M HCL in a beaker and mixed the contents well, adding dH20 to bring the total volume of the reaction to 10mL. We then microwaved the solution in a 750 watt oven for 120s. After removing the beaker from the microwave, we broke down the carbonized mass with a spatula. We added 15 mL of dH20 and continued to break down material in beaker. We then poured the liquid into a 50 mL falcon tube. To remove large particles, we centrifuged the solution at 15000 rpm for 20 min. We then removed the liquid supernatant with a 20 mL syringe and filtered with a 0.22 µm Millipore filter. To decrease the acidity of the nanoparticle solution, we changed the pH of the solution to 6 with KOH. To remove any large particles created by the pH change, we centrifuged the solution at 15000 rpm for 20 minutes, removed the supernatant with a 20 mL syringe, and filtered the solution with a 0.22 µm Millipore filter. We stored the particles at room temperature.

**1.8 Dextrin and Arginine**

We added 3g dextrin, 3g arginine and 0.1 or 0.05g bPEI 25k (made from the 100mg stock solution of bPEI listed in the notes) to 5 mL of 0.1 M HCL in a beaker and mixed the contents well, adding dH20 to bring the total volume of the reaction to 10mL. We then microwaved the solution in a 750 watt oven for 120s. A reddish mass will form. After removing the beaker from the microwave, we broke down the carbonized mass with a spatula. We added 20 mL of dH20 and continued to break down material in beaker. We then poured the liquid into a 50 mL falcon tube. To remove large particles, we centrifuged the solution at 15000 rpm for 20 min. We then removed the liquid supernatant with a 20 mL syringe and filtered with a 0.22 µm Millipore filter. To decrease the acidity of the nanoparticle solution, we changed the pH of the solution to 6 with KOH. To remove any large particles created by the pH change, we centrifuged the solution at 15000 rpm for 20 minutes, removed the supernatant with a 20 mL syringe, and filtered the solution with a 0.22 µm Millipore filter. We stored the particles at room temperature.

**1.9 Citric acid, Arginine, and Lysine**

We added 3g citric acid, 1.5g arginine, 1.5g lysine and 0.1 or 0.05g bPEI 25k (made from the 100mg stock solution of bPEI listed in the notes) to 5 mL of 0.1 M HCL in a beaker and mixed the contents well, adding dH20 to bring the total volume of the reaction to 10mL. We then microwaved the solution in a 750 watt oven for 120s. A yellow mass will form. After removing the beaker from the microwave, we broke down the carbonized mass with a spatula. We added 15 mL of dH20 and continued to break down material in beaker. We then poured the liquid into a 50 mL falcon tube. To remove large particles, we centrifuged the solution at 15000 rpm for 20 min. We then removed the liquid supernatant with a 20 mL syringe and filtered with a 0.22 µm Millipore filter. To decrease the acidity of the nanoparticle solution, we changed the pH of the solution to 6 with KOH. To remove any large particles created by the pH change, we centrifuged the solution at 15000 rpm for 20 minutes, removed the supernatant with a 20 mL syringe, and filtered the solution with a 0.22 µm Millipore filter. We stored the particles at room temperature.

**Chloroplast-targeting**

**1.10 Citric acid and Threonine**

We added 3g citric acid, 3g threonine and 0.1 or 0.05g bPEI 25k (made from the 100mg stock solution of bPEI listed in the notes) to 5 mL of 0.1 M HCL in a beaker and mixed the contents well, adding dH20 to bring the total volume of the reaction to 10mL. We then microwaved the solution in a 750 watt oven for 120s. After removing the beaker from the microwave, we broke down the carbonized mass with a spatula. We added 15 mL of dH20 and continued to break down material in beaker. We then poured the liquid into a 50 mL falcon tube. To remove large particles, we centrifuged the solution at 15000 rpm for 20 min. We then removed the liquid supernatant with a 20 mL syringe and filtered with a 0.22 µm Millipore filter. To decrease the acidity of the nanoparticle solution, we changed the pH of the solution to 6 with KOH. To remove any large particles created by the pH change, we centrifuged the solution at 15000 rpm for 20 minutes, removed the supernatant with a 20 mL syringe, and filtered the solution with a 0.22 µm Millipore filter. We stored the particles at room temperature.

**1.11 Citric acid and Glycine**

We added 3g citric acid, 3g glycine and 0.1 or 0.05g bPEI 25k (made from the 100mg stock solution of bPEI listed in the notes) to 5 mL of 0.1 M HCL in a beaker and mixed the contents well, adding dH20 to bring the total volume of the reaction to 10mL. We then microwaved the solution in a 750 watt oven for 120s. A glassy black mass will form. After removing the beaker from the microwave, we broke down the carbonized mass with a spatula. We added 20 mL of dH20 and continued to break down material in beaker. We then poured the liquid into a 50 mL falcon tube. To remove large particles, we centrifuged the solution at 15000 rpm for 20 min. We then removed the liquid supernatant with a 20 mL syringe and filtered with a 0.22 µm Millipore filter. To decrease the acidity of the nanoparticle solution, we changed the pH of the solution to 6 with KOH. To remove any large particles created by the pH change, we centrifuged the solution at 15000 rpm for 20 minutes, removed the supernatant with a 20 mL syringe, and filtered the solution with a 0.22 µm Millipore filter. We stored the particles at room temperature.

**Mitochondria-targeting**

**1.12 Citric acid, Arginine, Lysine, Serine**

We added 3g citric acid and 1.5g arginine, 1g serine, 1.5g lysine, and 0.1 or 0.05g bPEI 25k (made from the 100mg stock solution of bPEI listed in the notes) to 5 mL of 0.1 M HCL in a beaker and mixed the contents well, adding dH20 to bring the total volume of the reaction to 10mL. We then microwaved the solution in a 750 watt oven for 90s. After removing the beaker from the microwave, we broke down the carbonized mass with a spatula. We added 15 mL of dH20 and continued to break down material in beaker. We then poured the liquid into a 50 mL falcon tube. To remove large particles, we centrifuged the solution at 15000 rpm for 20 min. We then removed the liquid supernatant with a 20 mL syringe and filtered with a 0.22 µm Millipore filter. To decrease the acidity of the nanoparticle solution, we changed the pH of the solution to 6 with KOH. To remove any large particles created by the pH change, we centrifuged the solution at 15000 rpm for 20 minutes, removed the supernatant with a 20 mL syringe, and filtered the solution with a 0.22 µm Millipore filter. We stored the particles at room temperature.

**1.13 Citric acid and Serine (also go to nucleus)**

We added 3g citric acid, 3g serine and 0.1 or 0.05g bPEI 25k (made from the 100mg stock solution of bPEI listed in the notes) to 5 mL of 0.1 M HCL in a beaker and mixed the contents well, adding dH20 to bring the total volume of the reaction to 10mL. We then microwaved the solution in a 750 watt oven for 120s. A caramel-color mass will form. After removing the beaker from the microwave, we broke down the carbonized mass with a spatula. We added 15 mL of dH20 and continued to break down material in beaker. We then poured the liquid into a 50 mL falcon tube. To remove large particles, we centrifuged the solution at 15000 rpm for 20 min. We then removed the liquid supernatant with a 20 mL syringe and filtered with a 0.22 µm Millipore filter. To decrease the acidity of the nanoparticle solution, we changed the pH of the solution to 6 with KOH. To remove any large particles created by the pH change, we centrifuged the solution at 15000 rpm for 20 minutes, removed the supernatant with a 20 mL syringe, and filtered the solution with a 0.22 µm Millipore filter. We stored the particles at room temperature.

**1.14 Citric acid, Arginine, Lysine, and Valine**

We added 3g citric acid and 1g Arginine, 1g Lysine, 1g Valine and 0.1 or 0.05g bPEI 25k (made from the 100mg stock solution of bPEI listed in the notes) to 5 mL of 0.1 M HCL in a beaker and mixed the contents well, adding dH20 to bring the total volume of the reaction to 10mL. We then microwaved the solution in a 750 watt oven for 120s. A clear yellow, sticky mass will form. After removing the beaker from the microwave, we broke down the carbonized mass with a spatula. We added 15 mL of dH20 and continued to break down material in beaker. We then poured the liquid into a 50 mL falcon tube. To remove large particles, we centrifuged the solution at 15000 rpm for 20 min. We then removed the liquid supernatant with a 20 mL syringe and filtered with a 0.22 µm Millipore filter. To decrease the acidity of the nanoparticle solution, we changed the pH of the solution to 6 with KOH. To remove any large particles created by the pH change, we centrifuged the solution at 15000 rpm for 20 minutes, removed the supernatant with a 20 mL syringe, and filtered the solution with a 0.22 µm Millipore filter. We stored the particles at room temperature.

**Notes:**

1. For all the particles coated with bPEI, make a stock solution of bPEI25k (100mg/mL) as follows: add 5g bPEI 25k to 50mL dH20, stir with a magnetic stirrer, and microwave on high ca. 60 seconds until solution

starts to boil. Remove beaker and allow to cool to RT. bPEI is now soluble. If you do not follow these steps the bPEI will form large (ca. 1-5um) particles which causes DNA to condense immediately within 10 minutes and cells will not be transformed. Fukomoto et al. 2010 suggest you need to store PEI solutions in HCL. However, we have used stock bPEI made up with dH20 stored for 1-3 months without noticing any loss in transfection ability.

2. Transformation efficiency depends on bPEI concentration and organism. We have used three concentrations of bPEI for successful transformations- 0.2, 0.1 or 0.05g bPEI. Particles with a citric acid core are most effective with 0.2 or 0.1 bPEI; particles with a glucose or dextrin core are most effective when coated with 0.1 or 0.05 g bPEI. Plants and diatoms can handle higher levels of bPEI; algae are more sensitive and using 0.1 g bPEI works best.

2. Purification of nanoparticles

To purify nanoparticles of residual reagents, we used G10 and G100 Sephadex columns. To make the columns, we placed 1g Sephadex G10 or G100 into a 50mL Duran bottle. We added 30 mL H20 and placed the bottle in a 90 ° C hot water bath for 1h. After one hour, we poured out liquid at the top of the gel and allowed the gel to cool. 1 g of Sephadex made ca. 17-20 mL gel. We then took a glass HPLC column with a funnel and filter attached and added Sephadex gel to desired column height (1/2 to ¾ full). After filling the column with gel, we allowed the slurry to run out of the bottom. We then filled the column with equilibration buffer (for CNDs, this is dH20) three-times the column volume and discarded the run through. We took 1mL of the CNDs and washed them through a gel column with 3-5mL of dH20 (catching the run through in 500ul fractions) until the run through was clear (indicating no particles). This process diluted the particles, separated particles based on size, and presumably removed any residual noncarbonized particles. However, after comparing purified and non-purified particles, we did not notice any difference in cellular uptake or transformation.

3.Cell culture and growth conditions

*Arabidopsis thaliana* was grown on ½ MS media for 7-10 days under growth room conditions (continuous light, $23^0$ C). *Nicotiana benthamiana* plants were grown under greenhouse conditions (natural light-dark cycle at $25\text{-}27^0$ C) and infiltration experiments were performed when plants were 4-8 weeks old. *Selaginella kraussiana* was grown in a growth chamber (with a preset 16hour light/ 8 hour dark cycle at $23^0$ C); leaflets with rhizoids attached were cut from the mother plant and used in experiments. *Pseudomonas flourescens* was cultured on Mueller-Hinton Agar from frozen stock and incubated at 37° C for one week for biofilms to form. *Physarum polysephalum* was grown on 2% water agar and fed with oats, sub-cultured onto fresh plates every three days, and grown at 22° C. Fungi were grown from subcultures of hyphae (or spores) on 2% malt agar and grown at 22° C. *Penium margaritaceum* and *Spyrogira* sp. were grown in 50 mL flasks filled with Woods Hole media supplemented with vitamins; cultures were subcultured every 30 days and kept in a 16 hr light/dark cycle. *Thalassiosira oceanica, Synecococcus* sp., *Pleurochrysis certerae,* and *Gephyrocapsa oceanica* were received from Ros Rickaby's lab (Department of Earth Sciences, University of Oxford) in 15 mL Falcon tubes.

4. Nanoparticle uptake experiments

Nanoparticles were either added to agar plates or liquid growth media.  20 ul of p-CNDs was added to 100 mm plates filled with ½ MS and allowed to dry flat for 1 h before *A. thaliana* seedlings were added. Plates were divided in half so that the roots were exposed to the p-CNDs and the hypocotyl and leaves were not. We infiltrated *N. benthamiana* leaves with 100 ul of p-CNDs. *P. polycephalum* and *T. reesei* were grown on cellophane laid over 100 mm Petri dishes filled with malt agar for 24 h to form circular networks; these networks were then moved to split plates filled with 2 % water agar where one side was covered with 20 ul of CNDs and the other side was not covered with any treatment (control). Images were taken of the networks on the control side of the plate. Additional uptake experiments were performed with *P. polycephalum* by placing an oat flake inoculated with plasmodium on a 1 $cm^2$ piece of parafilm. After the physarum migrated off the parafilm and began to form a network (e.g. in 6-12 h), we added 5 µl of CND solution to the oatflake. We began imaging cultures after 30-60 min. to allow *P. polycephalum* to being breaking down the inoculated oatflake. We imaged nanoparticles flowing through the network at regular intervals of 1 cm until the network reached the edge of the plate. To check for cell uptake and toxicity of CNDs in algae, diatoms, and coccolithophores we added 15 ul of CNDs to 100 ul of stock cultures in 2 mL Eppendorf tubes. For uptake experiments, organisms were imaged at specified intervals from 30 minutes to 72 hours.

5. Transformation

The following plasmids were used in transformation experiments: pVenus NLS, pMCherry NLS, pGFP HDEL, and p35S ubi-GUS. Plasmids were received as a gift from Dr. Laura Moody (Department of Plant Sciences, University of Oxford). We inoculated 5 ul of CND-pDNA on 7-day-old *A. thaliana* roots and infiltrated *N. benthiana* leaves with 100 ul of CND-pDNA; plants were then returned to their growth chambers. To transform algae and diatoms, we added 15 ul of CND-pDNA to 100 ul of stock cultures in 2 mL Eppendorf tubes. To transform *Selaginella*, individual leaflets with rhizoids attached were placed in 2 mL Eppendorf tubes filled with 50 ul dH20 and 100 ul of CND-pDNA. All organisms were and imaged 12-72h and 8 days after inoculation. Transformation was usually seen 18h after exposure to CND-pDNA complexes. Organisms were exposed to the following treatments: control (no treatment), bPEI, bPEI + pDNA, CND, CND + pDNA. The above experiments were repeated with the following variables changed to optimize for transformation: pH of the CNDs (pH 1, 2, 4, 6); CND-pDNA complex time (10min, 15min, 20min, 30min, 60min, 24h); pDNA concentration (1ng, 100ng, 300ng, and 1ug/ul); and CND to pDNA ration (1:1, 1:2, 2:1).

6. Microscopy

Confocal images were taken using a Zeiss microscope using excitation 405/488/535/543nm. N. benthamiana and A. thaliana were mounted in perfusion chambers flooded with 500 ul Perfludecalin to remove air bubbles. A Leica dissecting microscope with removable gfp and yfp filters was used to image *P. polycephalum*, *T. reesei* and *Selaginella*.

**SI Table 1: Cellular localization of amino-acid based nanoparticles**

| Organelle | Composition | Organisms tested in |
|---|---|---|
| **Nucleus** | Citric acid and Lysine | Plants |
| | Citric acid and Arginine | Plants, algae |
| | Glucose and Arginine | Plants, algae |
| | Dextrin and Arginine | Plants |
| **Chloroplast** | Citric acid and Threonine | Plants, algae, diatoms, cyanobacteria |
| | Citric acid and Glycine | Plants |
| **Mitochondria** | Citric acid, Arginine, Lysine, Serine | Plants |
| | Citric acid and Serine (also go to nucleus) | Plants |
| | Citric acid, Arginine, Lysine, and Valine | Plants |
| **Cytoplasm** | Citric acid, Arginine, and Lysine | Plants |

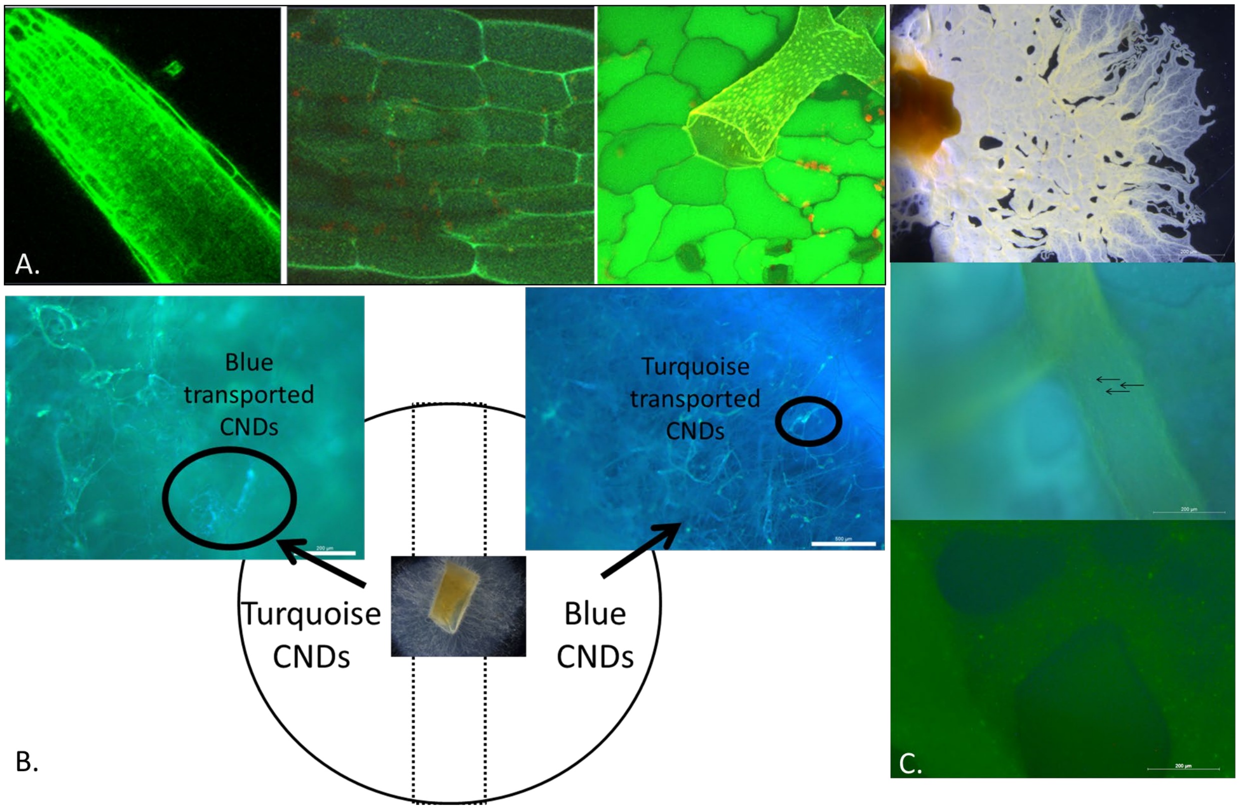


**SI Fig. 1. Long distance transport of carbon nanoparticles in living systems**. A. CND uptake by *A. thaliana* roots (left pane) and transport to hypocotyl (center pane) and leaf epidermis/trichomes (right pane). B. Transport of CNDs across hyphal networks by *T. reesei*. C. Transport of CNDs in *Physarum* plasmodium network (top pane) from agar coated with CNDs to non-coated area (center pane) and from a single central oat flake inoculated with CNDs (bottom pane). For all experimental designs and imaging, see SI Methods.

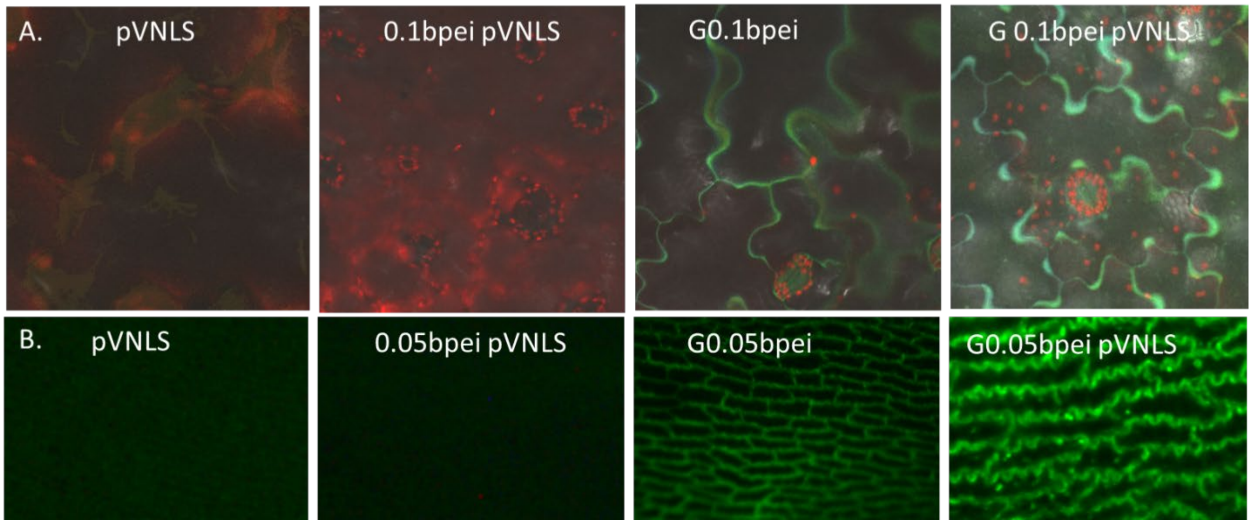


**SI Fig. 2: Control studies demonstrating CND-bPEI-pDNA complexes are needed for successful transformation in *N. benthamiana* (A) and *Selaginella* (B).** In the control images, only plant autoflorescence is seen when exposed to either the plasmid or the plasmid complexed with bPEI. Cells are outlined when exposed to the nanoparticle without a plasmid attached. Nuclei are fluorescent when exposed to a carbon nanoparticle with a VNLS plasmid attached. Abbreviations explained: pVNLS-plasmid encoding a Venus yellow fluorescent protein with a nuclear localization signal; 0.1/0.05 bpei pVNLS- different concentrations of bPEI and the pVNLS plasmid; G0.1/0.05bpei-a glucose based carbon nanoparticle coated with either 0.1 or 0.05 g bPEI; G0.1/0.05bpei pVNLS-glucose carbon nanoparticle coated with bPEI with a VNLS plasmid attached. Confocal images (A) are composed of superimposed z-sections. YFP was imaged with 535 excitation on confocal and are shown as yellow/green in the images; chloroplast autoflorescence is red. The images in B were taken with a Leica light microscope with a YFP filter.
